# The Impact of Protein Dynamics on Residue-Residue Coevolution and Contact Prediction

**DOI:** 10.1101/2022.10.16.512436

**Authors:** Alexander Fung, Antoine Koehl, Milind Jagota, Yun S. Song

## Abstract

The need to maintain protein structure constrains evolution at the sequence level, and patterns of coevolution in homologous protein sequences can be used to predict their 3D structures with high accuracy. Our understanding of the relationship between protein structure and evolution has traditionally been benchmarked by computational models’ ability to predict contacts from a single representative, experimentally determined structure per protein family. However, proteins *in vivo* are highly dynamic and can adopt multiple functionally relevant conformations. Here we demonstrate that interactions that stabilize alternate conformations, as well those that mediate conformational changes, impose an underappreciated but significant set of evolutionary constraints. We analyze the extent of these constraints over 56 paralogous G protein coupled receptors (GPCRs), *β*-arrestin and the human SARS-CoV2 receptor ACE2. Specifically, we observe that contacts uniquely found in molecular dynamics (MD) simulation data and alternate-conformation crystal structures are successfully predicted by unsupervised language models. In GPCRs, adding these contacts as positives increases the percentage of top contacts classified as true positives, as predicted by a state-of-the-art language model, from 69% to 87%. Our results show that protein dynamics impose constraints on molecular evolution and demonstrate the ability of unsupervised language models to measure these constraints.

## 1 Background

In recent years, the exponential decrease in sequencing costs has led to rapid growths of protein sequence databases that has vastly outstripped the rate of experimental structure determination and functional annotation. This discrepancy has led to significant interest in the development of unsupervised learning methods that can extract biologically meaningful information from sequences alone. In particular, many techniques focus on protein families – sets of homologous sequences that share a common underlying fold and function while retaining as little as 15% sequence identity between individual sequences. These protein families are an evolutionary record of the need to balance exploration and conservation, with maintenance of structure and function imposing strong restraints on sequence variation within a family. In order to maintain interactions important for function, residues that are in close 3D spatial proximity (even if distant in sequence) show strong signals of co-evolution [1–3]. The breakthrough success of AlphaFold2 and other methods in predicting protein structure from sequence depends heavily on learning patterns of evolutionary variation, either from explicit sequence alignments [4, 5] or implicit representations of evolutionary patterns [6].

While AlphaFold2 and related work have greatly advanced our understanding of how protein structure influences sequence evolution, this understanding is centered around a static view of protein structure. AlphaFold2 is trained on individual protein structures that are mostly resolved by X-ray crystallography, and unsupervised methods that preceded AlphaFold2 have mostly been evaluated on these same static structures [7, 8]. In contrast, proteins *in vivo* often depend on significant dynamical changes to carry out their functions. Many proteins, for example, adopt alternate conformations in complex with another protein or small molecule binder. AlphaFold2 has been shown to possess some implicit knowledge of protein dynamics [9, 10], but there is no clear way to extract such information at high resolution for an arbitrary protein of interest. Training a deep learning model to predict dynamics of a protein based on sequence alone remains out of reach due to the scarcity of experimental dynamics measurements and the computational intensiveness of molecular dynamics simulations [11].

We postulated that protein dynamics might influence sequence variation in evolutionarily related proteins beyond the limited role which has been previously established [12, 13]. We collected protein structures for three different families representing a larger fraction of each conformational landscape than is typically analyzed. Specifically, we gathered experimental structures of 56 paralogous G protein coupled receptors (GPCRs), *β*-arrestin, and the human SARS-CoV2 receptor ACE2 in multiple conformational states, as well as microsecond timescale molecular dynamics launched from those states. Then, as illustrated in **Figure 1**, we measured the ability of protein language models to predict interactions (residue-residue contacts) that only occur in alternate dynamical states. Rather than using AlphaFold2, which has been trained to predict stable crystal structures, we chose to use unsupervised models of protein sequence variation, which capture all types of co-evolutionary constraints without bias. We primarily used the protein language model MSA Transformer, which is the highest-performing unsupervised method for the prediction of static structure protein contacts [14]. We found that dynamics-only interactions are predicted by MSA Transformer with higher strength than can be explained based on single static structures. Since these models are trained on protein sequence variability data, our results demonstrate that dynamics constrain sequence evolution and illustrate the importance of understanding the full conformational landscapes of proteins. Our results also show that assessments of unsupervised language models of sequence variation based on static structure *underestimate* model performance; for GPCRs, including dynamic interactions in an evaluation of MSA Transformer means that it identifies interactions with a precision of 87% compared to 69% using static interactions alone, when the number of predicted contacts is equal to the length of the protein.

**Fig. 1.**
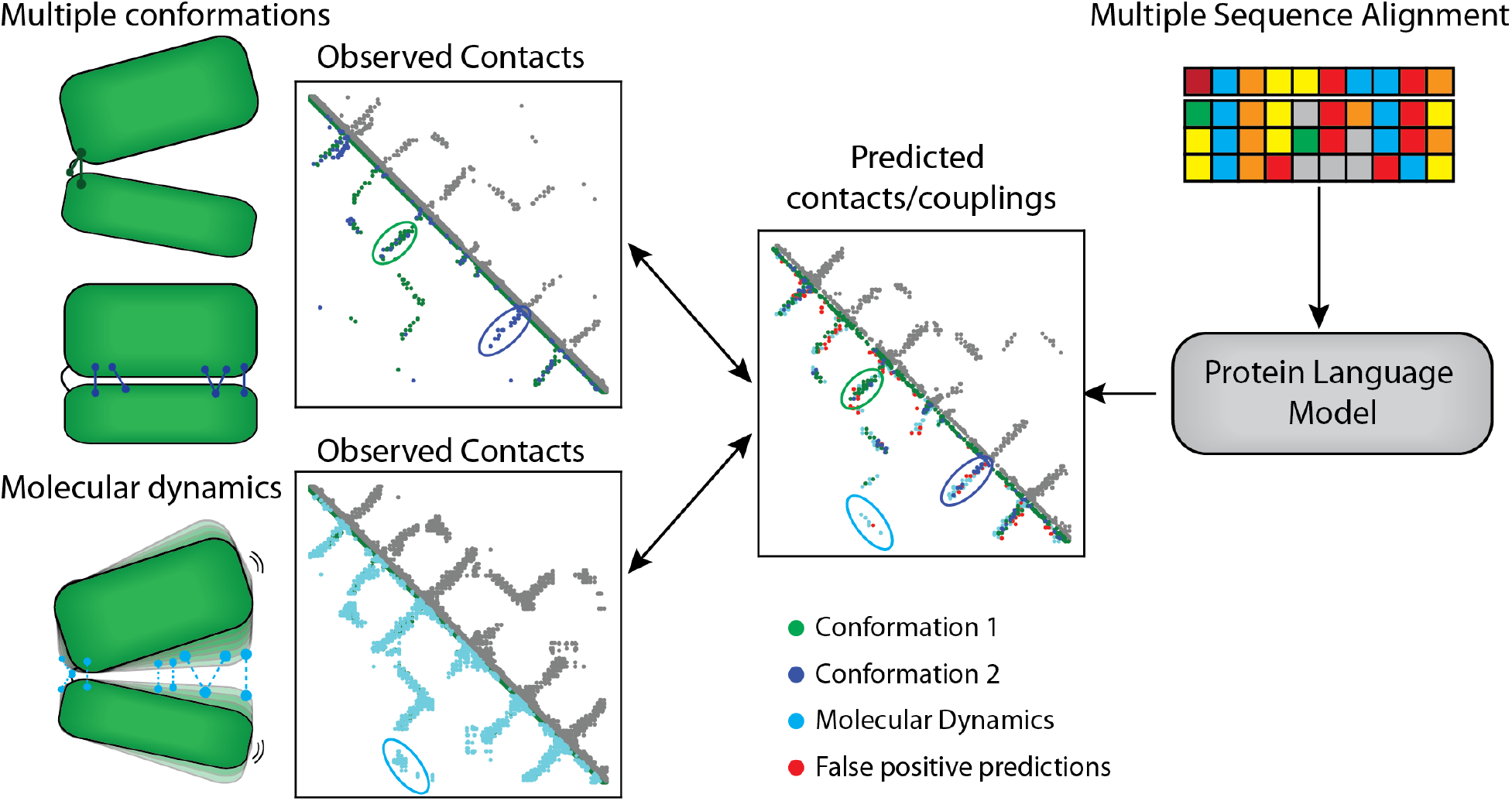
Evaluating contact prediction across proteins’ conformational landscapes. Proteins adopt different conformations and transient dynamical states, each with unique sets of contact patterns (cartoon-left). Protein language models, which are trained only on evolutionary protein sequence data, can be used to predict contacts (right). We observe that protein language models are able to predict contacts unique to alternate conformations (top left) as well as those unique to transient dynamical states (bottom left). These effects show the impact of protein dynamics on sequence coevolution.

## 2 Methods

### 2.1 Datasets

We studied contact maps across the conformational landscape for 56 paralogous GPCRs, as well as for *β*-arrestin and ACE2. For each protein, we generated ground truth contact maps using the following data: a representative experimental structure, molecular dynamics (MD) simulation trajectory data, and the experimental structure used to initialize the simulation. For GPCRs, representative structures were obtained from the PDBRenum server [15], which renumbers residues in Protein Data Bank (PDB) [16] files to exactly match their UniProt [17] sequence for ease of downstream processing. MD trajectory data and corresponding initialization structures were obtained from GPCRmd [18]. For *β*-arrestin, representative structures were obtained from the PDB, while trajectories and initialization structures were obtained from Latorraca *et al*. [19]. ACE2 trajectories and initialization structures were obtained from the D.E. Shaw Research SARS-CoV-2 technical data repository [20].

### 2.2 Contact Extraction and Prediction

We extracted both static and dynamic contacts with the GetContacts software package [21]. GetContacts identifies direct inter-residue contacts based on a combination of contributing atom types and interaction geometry. Additionally, we calculated C_*β*_-distance-based contact maps which classify residues as being “in contact” if the distance between their C_*β*_ atoms is less than 8.5Å [22]. For each protein, we identified the subset of residues that were present in the structure files for all conformations of that protein, and truncated the extracted contact maps to only include interactions between those residues. We were able to compute contact maps for 56 out of 68 total GPCRs in the GPCRmd dataset, as well as bovine *β*-arrestin and human ACE2, which are each a single protein.

Predicted contacts were generated using pretrained MSA Transformer [14] and ESM-1b models [23]. We extracted the UniProt sequences corresponding to our experimental structure data and generated MSAs for these sequences using HHblits [24] to search against the UniRef database. MSA Transformer inference was run on the MSAs, while ESM-1b model inference was performed on single sequences directly. Transformer-based language models, like MSA Transformer and ESM-1b, build internal representations of the way different positions in their inputs relate to each other in the form of *L* × *L* attention matrices, where *L* is the input protein sequence length [25]. Previous work has shown that a sparse combination of these attention matrices can accurately predict protein contact maps [8, 26]. We evaluate the precision (defined as the percentage of predicted contacts classified as true positives) on various fractions of the top *L* attention activations in order to account for the roughly linear scaling of contacts in relation to protein length *L* [27, 28].

The precision metrics we chose were motivated by the standard baseline of a single static structure, which we denote as the *primary* structure, and we analyzed precision increases resulting from adding contacts from either alternate conformation (*secondary*) static structures or dynamics to the set of positives. Therefore, each metric was computed on a per-conformation basis. We report precision on the following: C_*β*_-distance-based contacts, GetContacts statics, aggregate contact maps consisting of statics from multiple conformations, GetContacts dynamics, and combinations of the aforementioned. For each metric, we computed precision on top-*L*, top-*L*/2, and top-*L*/5 contacts across short-range (6 ≤ sep ≤ 12), medium-range (12 ≤ sep ≤ 24) and long-range (24 ≤ sep) contacts, where “sep” denotes the inter-residue distance in 1D sequence.

### 2.3 Control Precision Values

We evaluated the precision of MSA Transformer for contact prediction using various definitions of positives resulting in different numbers of true positives. Precision values assessed over different numbers of positives are not fully controlled. In order to test the significance of observed precision increases, we calculated a control precision for each set of positive contacts, in which we sampled an equal number of contacts with the same distribution of reference static structure distances and instead defined those as positives. This control measures how strongly a set of contacts is predicted compared to an alternative based purely on distances in one representative structure. Concretely, we used the following scheme to randomize contact maps: bin a subset of non-diagonal residue pairs by C_*β*_-C_*β*_ distance into 21 bins according to the following divisions (in Å): {0, 2.5, 3.0, 3.5, 4.0, 4.5, 5.0, 5.5, 6.0, 6.5, 7.0, 7.5, 8.0, 8.5, 9.0, 10.0, 12.0, 18.0, 24.0, 36.0, 50.0, ∞}, then randomly permute contacts within each bin. For primary statics and dynamics metrics, all contacts were permuted. All other metrics involved adding new positives to a primary statics contact set, and for these metrics all contacts not in the primary statics were permuted.

## 3 Results

### 3.1 MSA Transformer Predicts Dynamic Contacts

MSA Transformer is an unsupervised protein language model, trained to understand sequence variability in protein multiple sequence alignments. From databases of protein sequences alone, MSA Transformer learns which pairs of positions in a particular protein have significant covariance and stores this information in attention maps. These attention maps correlate strongly with static structure contacts.

We find that MSA Transformer attention maps capture contacts across the conformational landscapes of proteins, and not just in single static structures. We collected contacts from 56 GPCRs, of which 42 had structures in both inactive and active conformations, as well as from ACE2 and *β*-arrestin. We computed the precision of predicted MSA Transformer contacts using various sources of contacts as positives: a representative static structure, additional structures in an alternate conformations, or a molecular dynamics simulation (**Table 1)**. We also performed all the same evaluations using ESM-1b, with similar trends but lower accuracy for ESM-1b on all metrics. We report these evaluations and expanded results in **Supplementary Table S1**.

**Table 1.** Average precision across all proteins for MSA Transformer predictions. “C_*β*_-distance” refers to contacts extracted from an experimental structure based on a C_*β*_-C_*β*_ distance threshold of 8.5Å, while “Static” refers to contacts extracted from an experimental structure via GetContacts. “Agg” refers to combining contacts from multiple conformations. “Dyn” refers to adding contacts from molecular dynamics. Unparenthesized numbers are true precisions, and parenthesized (GPCRs only) are baseline precisions computed via the equidistant C_*β*_-C_*β*_ control. Adding new positives from dynamical information consistently increases precision beyond the control, indicating that contacts from dynamics are significantly predicted. Each precision value is the mean precision of all conformations of a protein. Each GPCR value is the mean over the aforementioned mean precision for all GPCRs. (14 GPCRs had only one static conformation, so aggregate statistics were omitted.) Overall GPCR precision was computed for the top-*L*, top-*L*/2, and top-*L*/5 contacts. For short-range (6 ≤ sep ≤ 12), medium-range (12 ≤ sep ≤ 24) and long-range (24 ≤ sep) GPCR contacts, we computed precision over the subset of the top *L* predicted interactions with the corresponding separation.

| Metric | All GPCRs ($6 \leq \text{sep}$) | | | Short | Medium | Long | ACE2 | β-arrestin |
| --- | --- | --- | --- | --- | --- | --- | --- | --- |
| | $L$ | $L/2$ | $L/5$ | $L$ | $L$ | $L$ | $L$ | $L$ |
| $C_\beta$-distance | 0.69 (0.69) | 0.82 (0.82) | 0.91 (0.91) | 0.58 (0.58) | 0.77 (0.77) | 0.78 (0.78) | 0.62 | 0.62 |
| Static | 0.37 (0.33) | 0.47 (0.43) | 0.54 (0.50) | 0.30 (0.26) | 0.40 (0.38) | 0.42 (0.38) | 0.30 | 0.32 |
| Agg $C_\beta$-distance | 0.74 (0.72) | 0.86 (0.84) | 0.93 (0.92) | 0.63 (0.61) | 0.81 (0.80) | 0.83 (0.80) | 0.70 | 0.66 |
| Agg Static | 0.50 (0.47) | 0.62 (0.58) | 0.70 (0.67) | 0.41 (0.38) | 0.53 (0.51) | 0.57 (0.53) | 0.41 | 0.40 |
| Dyn | 0.81 (0.72) | 0.89 (0.82) | 0.93 (0.88) | 0.72 (0.64) | 0.81 (0.79) | 0.88 (0.78) | 0.80 | 0.62 |
| Agg $C_\beta$-distance + Dyn | 0.87 (0.80) | 0.94 (0.89) | 0.98 (0.94) | 0.78 (0.72) | 0.89 (0.87) | 0.93 (0.86) | 0.84 | 0.69 |
| Agg Static + Dyn | 0.84 (0.76) | 0.92 (0.85) | 0.96 (0.91) | 0.74 (0.67) | 0.84 (0.81) | 0.90 (0.82) | 0.80 | 0.63 |

Broadly, inclusion of alternate conformation and dynamic contacts as positives led to significant gains in apparent precision relative to control across all ranges and at all prediction levels. This gain was especially dramatic for the long-range contact subset (sep ≥ 24), as defined by contacts involving residues separated by 24 or more positions in 1D sequence. Such contacts are particularly suited to determining global structure, and novel interactions in this region are more likely to correspond to large domain conformational changes. For long-range contacts, we observe an increase of 15% in precision by including contacts from alternate crystal structures as positives, and an additional increase of 36% by further including contacts unique to molecular dynamics.

The addition of new positives to contact prediction can only increase precision. To determine whether increases in precision were meaningful, we computed a control precision for each measurement, as described in Section 2.3. This control was calculated as the precision of MSA Transformer over an equal number of positives, sampled randomly with a matching distribution of static structure distances. This control assesses whether an increase in precision can be explained by simply picking positives based on distance in one representative structure. All methods of adding positives from dynamical information increased precision beyond the control, indicating that contacts from dynamics are predicted by MSA Transformer at a level beyond what can be expected from a representative static structure.

The substantial improvement in precision given by dynamics over the C_*β*_-distance baseline indicated that many dynamic interactions have C_*β*_-C_*β*_ distances greater than 8.5A in the primary static structure. We analyzed distributions of C_*β*_-C_*β*_ interaction distances for various interaction types (**Figure 2)** and observed that dynamic contact pairs, and to a lesser extent contacts from alternate conformation static structures, are often distant in primary static structures and therefore would not be detected by distance threshold relaxation in C_*β*_-distance-based methods. We additionally examined distributions of sequence separation by contact type, observing similar trends (**Supplementary Figure S1)**.

**Fig. 2.**
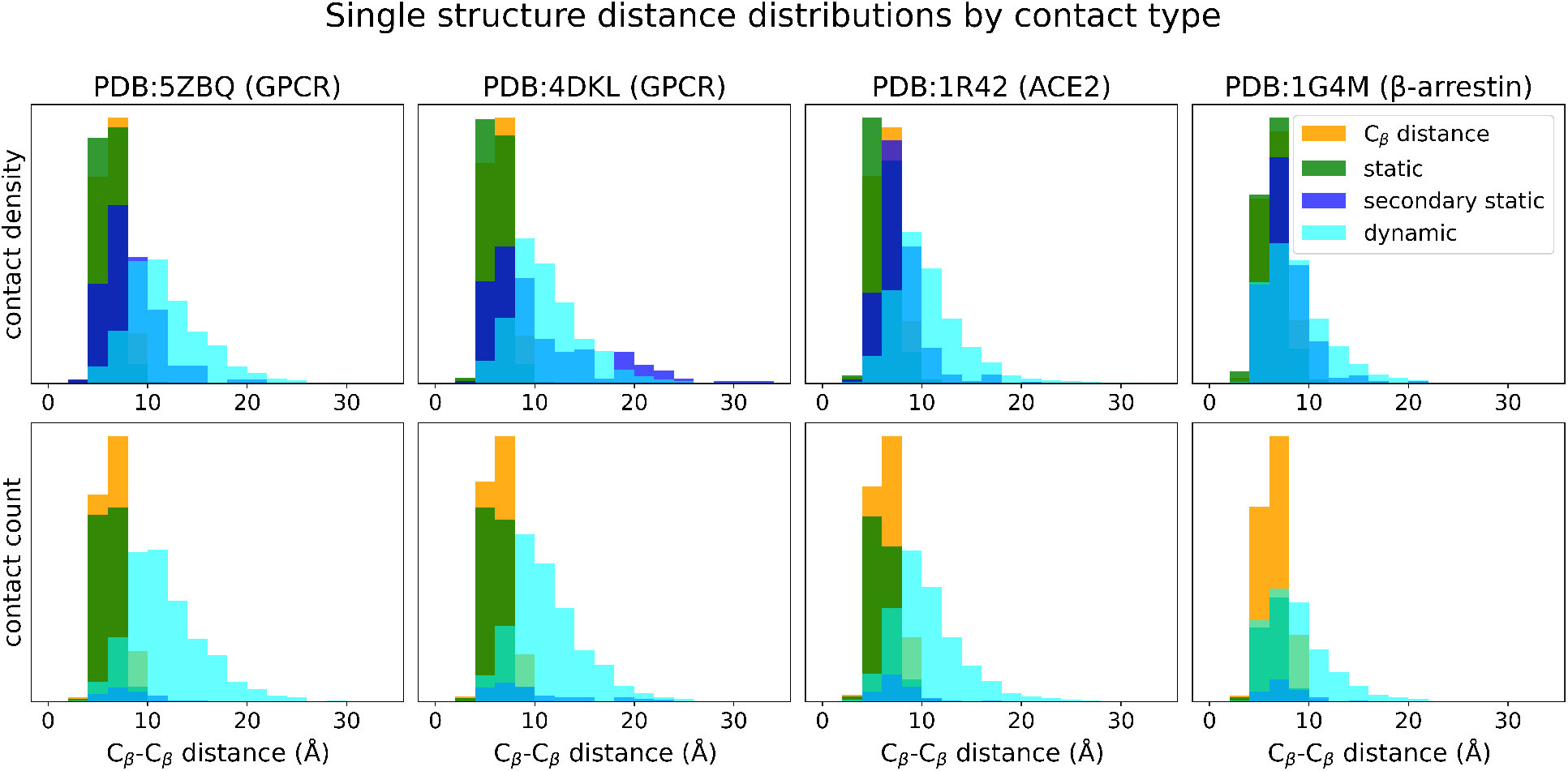
Distributions of residue-residue distance in representative static structure for unique contacts of each type. C_*β*_-distance and static refer to different ways to extract contacts from the representative static structure (C_*β*_-distance or GetContacts, respectively). Secondary static contacts are those contacts unique to an alternate conformation, and dynamic contacts are unique to molecular dynamics simulations. A large fraction of new contacts from dynamics and alternate conformations are distant in the representative conformation, meaning these contacts would be hard to predict from a single structure. Molecular dynamics introduces far more of these novel contacts in absolute number.

We report precision for both GetContacts statics and C_*β*_-distance-inferred static interactions. Classifying contacts based on pairwise C_*β*_ distances has been a mainstay in the field of contact prediction, beginning with its incorporation as a standard metric in Critical Assessment of protein Structure Prediction (CASP) challenges [29]. Compared to direct interaction-based contact extraction techniques, C_*β*_-distance-based methods have the advantages of computational simplicity, and are side chain-agnostic, making them more robust to inaccurately modeled side chains in lower-resolution structures. However, prior work has noted that such methods tend to inflate precision, and predominantly report on contact pairs that contribute little to structural interpretation [22]. This further motivated the equidistant C_*β*_ control, which directly compares GetContacts and C_*β*_-distance-based contact extraction methods on a per-contact basis. Across our GPCR data, we find that contacts extracted from a single static structure using GetContacts evaluate with higher top-*L* precision (37%) than control (33%), suggesting that they provide structural insights beyond what can be inferred by simply looking in the neighborhood of residues that are already near to one another. We henceforth refer to contacts extracted from static structures using GetContacts simply as “statics,” and focus the remaining analysis on the degree to which molecular dynamics and alternate conformation structures improve upon these “statics.”

### 3.2 Predicted Contact Probabilities Correlate with Dynamic Contact Dwell Time

Contacts that appear in molecular dynamics simulations generally do not persist over the whole simulated time. We examined whether MSA Transformer could predict the duration of contacts in molecular dynamics simulations. We restricted to contacts that are unique to molecular dynamics and then computed the dwell time fraction for each, defined as the fraction of the MD simulation time where the contact exists. We also restricted to contacts with sep ≥ 6. We then correlated these dwell times with the predicted probability of contact from MSA Transformer (**Figure 3)**. Across all protein families, there is significant correlation between dwell time fraction and predicted contact probability from MSA Transformer, with particularly high correlation in GPCRs. This result holds despite the fact that we have removed many of the strongest contacts by only using those which are unique to molecular dynamics. The magnitude and statistical significance of these correlations is robust to the sequence separation cutoff we employed (**Supplementary Figure S2)**. Typically, the contact probability is only used as a means of ranking contacts for selection at different thresholds as a function of sequence length (e.g., to find the “top” *L* contacts). By showing that predicted contact probability correlates with the energetics of an interaction, we provide a justification for more quantitative interpretation of the outputs of protein language models. In **Table 2**, we list the mean Spearman’s rank correlation *ρ* for each protein family.

**Fig. 3.**
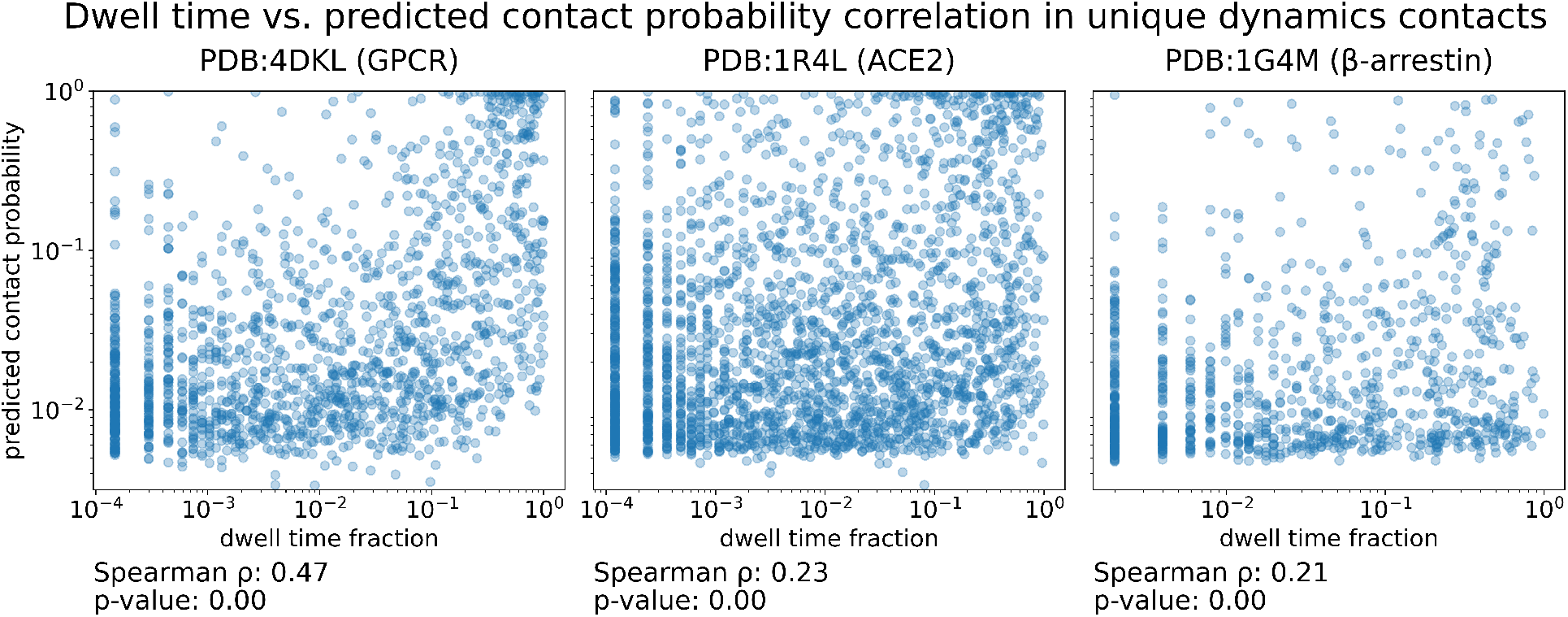
Prediction strength correlates with dwell time. We calculated dwell time fraction for all contacts unique to molecular dynamics with sequence separation ≥ 6 and correlated this with the predicted contact probability from MSA Transformer. We define dwell time fraction as the fraction of time that a particular contact is formed across a molecular dynamics trajectory. Contacts with higher predicted contact probability correlate with increased dwell time, even though we have removed many of the strongest contacts by only using those which are unique to molecular dynamics. Empirical plots of dwell time fraction against predicted contact probability are shown for a representative GPCR, the *μ*-opioid receptor, as well as ACE2 and *β*-arrestin. Note that axes are on a log scale.

**Table 2.**
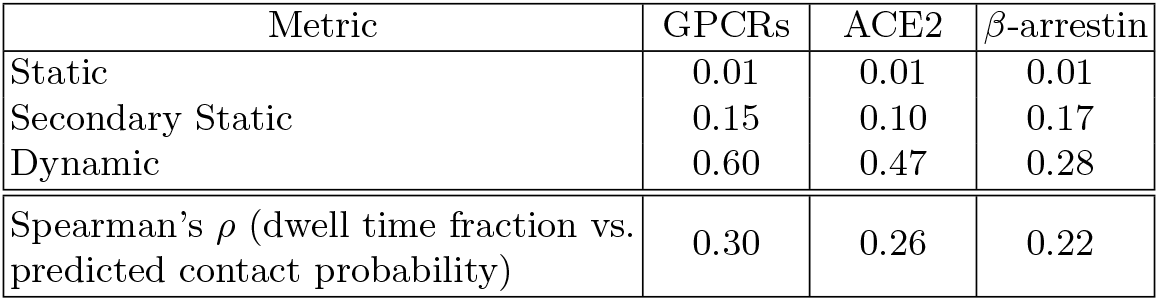
Fraction of of distant contacts and correlation between dwell time and prediction strength. Distant contacts are defined as residue pairs separated by least 10Å; contacts at this distance are out of range for reasonable C_*β*_-C_*β*_ thresholding schemes. Across all protein families, molecular dynamics provide a significant source of new interactions between residues originally too far apart – likely corresponding to real conformational changes. There are significant Spearman correlations between dwell time fraction and MSA Transformer predicted contact probability for unique dynamics in each protein family. (Many strongly predicted, high dwell time contacts are also present in primary static structures, so we omit them to avoid inflating correlation.) GPCR metrics are also the mean across all GPCRs.

### 3.3 Family-Specific Biological Insights

#### GPCRs

G protein-coupled receptors (GPCRs) are the largest family of cell surface receptors in eukaryotes, with 800 members in the human genome [30]. As a family, they share a common fold that is characterized by seven transmembrane (TM) *α* helices and are fully embedded in the cell membrane (**Figure 4**a). As cell surface receptors, GPCRs become activated by binding to ligands on the outside of the cell, at which point they can initialize intracellular signaling cascades that lead to changes in cellular behavior [31]. While the structural hallmark of active receptors is a large outward motion of TM helix 6 (**Figure 4**a), biophysical and molecular dynamics studies have shown that GPCR activation is not a simple “on-off switch”, but rather a consequence of increased local dynamics surrounding TM6 [32].

**Fig. 4.**
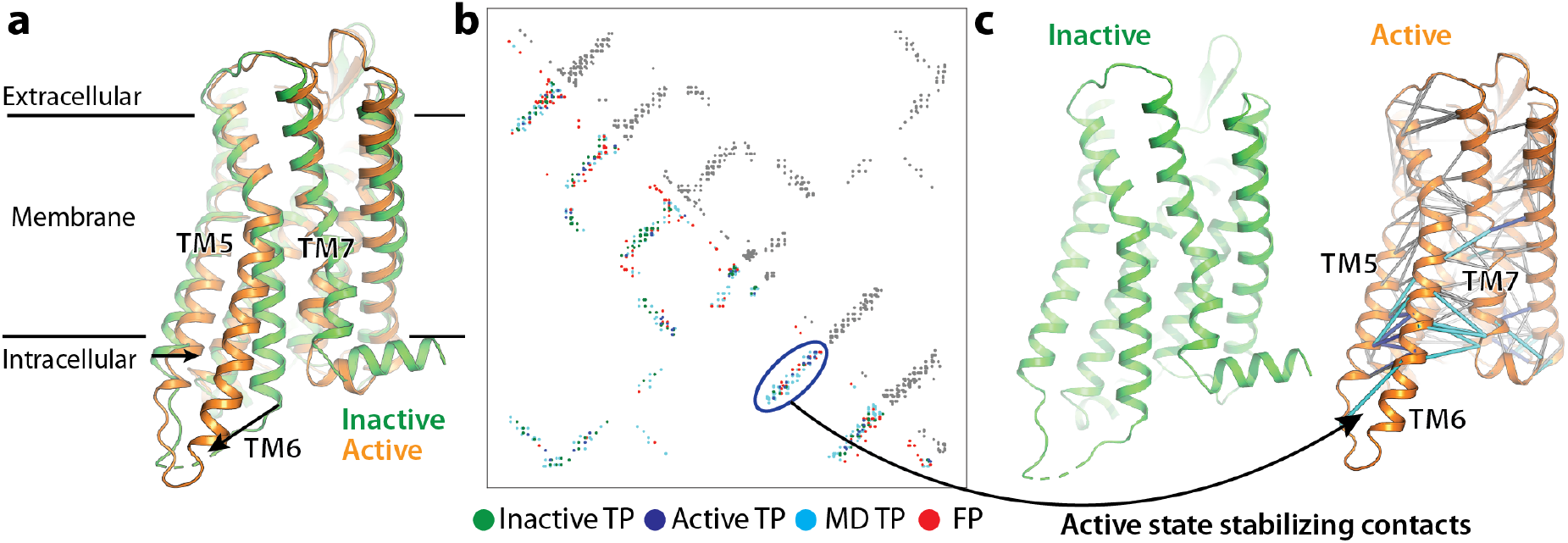
Prediction of GPCR active-state-stabilizing contacts. **(a)** Comparison of the *μ*-opioid receptor in inactive (green) and active (orange) states. **(b)** Top *L* MSA Transformer predicted contacts. In the upper triangular part of the matrix, all predictions are colored gray, whereas in the lower triangular part, predictions are colored by the state in which they are uniquely observed (green, blue, cyan) or false positives (red). **(c)** Correctly predicted contacts unique to active state of the *μ*-opioid receptor are shown as lines in the active state crystal structure (PDB:5C1M). Lines are colored blue if they correspond to residues that show large positional changes in crystal structures, and gray otherwise.

In particular, we frequently observe a cluster of high-confidence predicted contacts between TM5 and TM6 that uniquely arises when these transmembrane helices adopt their active-state conformations (**Figure 4**b,c – here shown for the *μ*-opioid receptor). This TM5-TM6 interface we highlight involves only a single contact pair from the inactive state structure, in stark contrast to the 10 present in the active state structure, and 49 in molecular dynamics. Predicted contacts from MSA Transformer in this region demonstrate impressive recall; they correctly identify 9 of the 10 active-state contacts and 12 of the 49 dynamical contacts (**Figure 4**b,c), showing high sensitivity to detect contact networks involved in important biological processes. (We also observe new interactions at the TM6-TM7 interface in molecular dynamics, which occur in states even more “inactive” than the initial state – thus highlighting the spectrum of TM6 motion.) Though the relative stability of the inactive state necessitates intracellular stabilizing proteins to obtain active-like receptor structures [33–35], we find that interactions contributing to, and stabilizing, this TM6 outward motion are also a strong source of signal in sequence variation, as 15 of the 60 new positive contacts originating from the active state structure cluster at the cytoplasmic ends of TM5 and TM6.

#### ACE2

The angiotensin-converting enzyme (ACE)-related carboxypeptidase, ACE2, is a membrane anchored enzyme that is involved in degradation of the blood pressure modulating peptide angiotensin, and perhaps more notably, is the human receptor for the SARS-CoV-2 virus [36]. Structurally, ACE2 is composed of two domains, an N-terminal metalloprotease domain (NTD) that is responsible for peptide cleavage, as well as a C-terminal collectrin-like domain (CTD). Binding of ACE2 inhibitors, and presumably the native substrate angiotensin, leads to a ~ 16° hinge bending motion of the two domains relative to each other that leads to a closing of the substrate cleft [37] (**Figure 5**b).

**Fig. 5.**
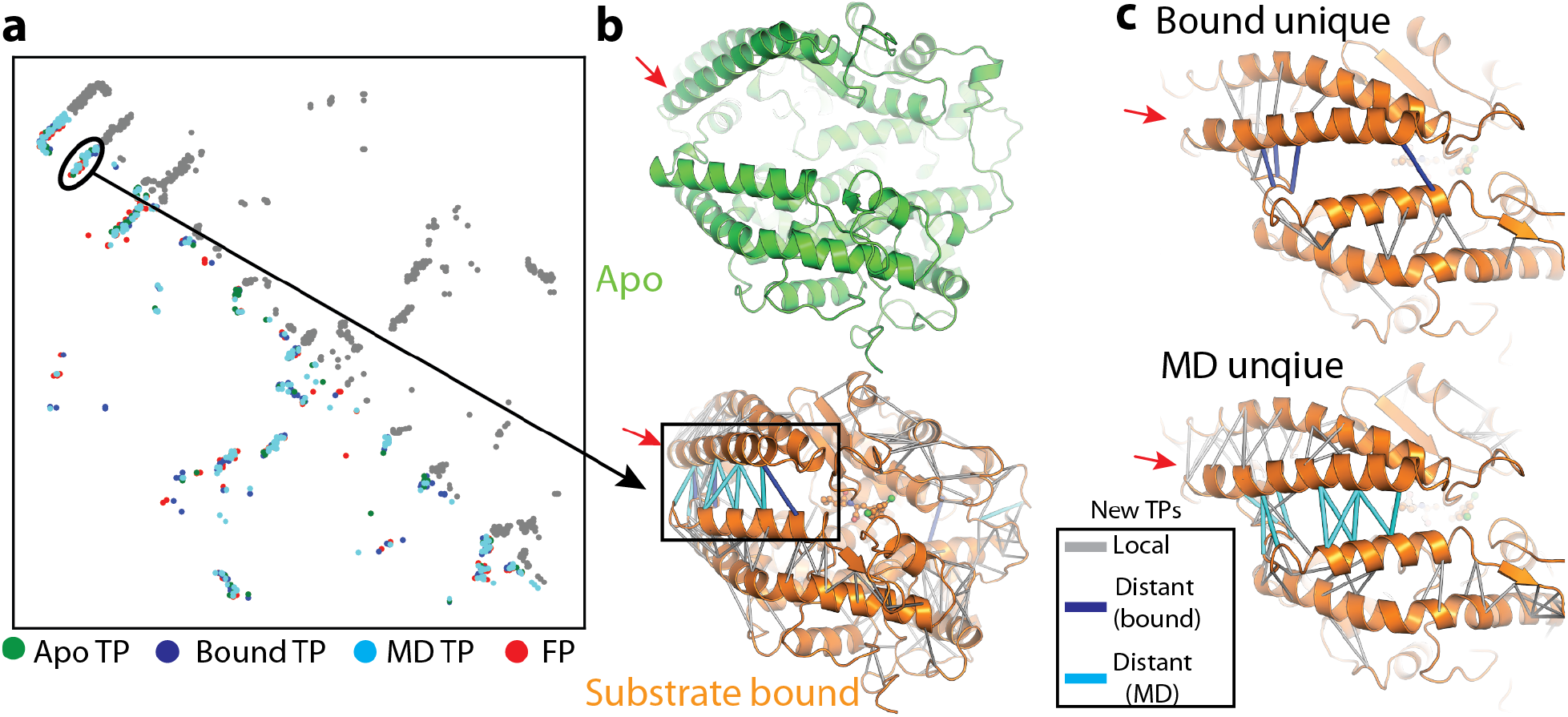
Language models correctly identify dynamic and state-specific contacts in ACE2. **(a)** Top *L* MSA Transformer predictions shown as gray circles (upper right), and colored (lower left) red if false positives, or by provenance. **(b)** Comparison of Apo (top, green) (PDB:1R42) and substrate-bound (bottom, orange) (PDB:1R4L) ACE2 showing the result of lid closure on a distal helix (red arrow). Correctly predicted contacts unique to the MD simulation or the inhibitor bound state are mapped onto the substrate bound complex and colored if they correspond to residues that show large positional changes upon substrate binding (cyan = from MD, blue = substrate bound), or gray if they report on relatively immobile regions. **(c)** Zoomed inset indicated in **b** showing top predicted contacts unique to the substrate bound state (top) or the MD simulation (bottom). Coloring is identical to **b**.

Compared to evaluating precision on the apo structure alone, inclusion of novel contacts from both the structure of substrate bound, as well as MD trajectories launched from the apo-state structure, lead to greatly increased precision metrics of contacts predicted by language models (**Figure 5**a). While the majority of new correctly predicted contacts involve local sets of interactions that likely could be captured by relaxing contact inclusion criteria (**Figure 5**b,c), we observe a series of contacts clustered between two apical helices (**Figure 5**a (circled), b&c (red arrows)) capturing the most significant motion between apo- and substrate-bound structures. In particular, these contacts are unique to the substrate bound conformation and as such, primarily report on lobe closure. More specifically, when focusing on the contact region between these two helices, we see that only a single residue pair is present in the apo-structure, compared to 9 and 50 from the substrate-bound and MD trajectory, respectively. Impressively, 6 (of 9 observed) substrate-bound and 13 (of 50 observed) MD simulation contacts account for all 19 new true positive contacts in this region, demonstrating high recall for a dynamically important region. The formation of contacts that stabilize both apo- and substrate-bound states are important for ACE2’s enzymatic activity; these sets of interactions are likely important drivers of coevolution.

#### *β*-Arrestin

*β*-arrestin is a soluble protein that is predominantly composed of *β* strands and is involved in modulation of GPCR signaling pathways. Canonically, *β*-arrestin becomes activated by binding to phosphorylated C-terminal tails of activated GPCRs, leading to desensitization and internalization of the receptor, as well as initiation of arrestin-specific signaling cascades [19]. Structurally, *β*-arrestin is an elongated protein composed of two folded domains (termed N-lobe and C-lobe) that are composed of contiguous sequence segments corresponding to the first and second halves of the sequence, respectively. These domains share a relatively restricted interface, with the majority of interdomain contacts arising from a sparse set of residues. A combination of molecular dynamics and biophysical work has shown that activation of *β*-arrestin predominantly involves fairly local, domain restricted, conformational changes, particularly in a series of apical loops. Indeed, we observed that incorporation of MD data and alternate conformations into the analysis of predicted contacts led to increased precision, but by a modest amount compared to the aforementioned improvement for GPCRs and ACE2 (**Table 2)**. This may be reflected by only ~28% of unique dynamic interactions in *β*-arrestin being separated by at least 10Å, in contrast to ~60% in GPCRs and ~47% in ACE2.

## 4 Discussion

We showed that dynamical changes of proteins constrain sequence evolution in ways that cannot be predicted from single static structures. In GPCRs, *β*-arrestin, and ACE2, residue-residue interactions that are unique to alternate conformational states and transient dynamical changes are accurately predicted by MSA Transformer, an unsupervised model of protein sequence variation. We relied on molecular dynamics simulations and crystal structures of alternate conformations for ground truth data about dynamical changes of proteins. In the future, larger amounts of molecular dynamics data could enable further investigation into the relationship between protein dynamics and sequence evolution. We are also excited about the rapid growth of Cryo-EM as a method of structure determination. Cryo-EM allows the capture of protein structures in more varied and natural structural states, as well as the direct inference of continuous variation [38–40], and could therefore provide a large additional source of information about protein dynamics. As both of these data sources grow, it will be interesting to consider integrating dynamics prediction into a supervised structure predictor such as AlphaFold2 [4].

MSA Transformer and other deep learning methods of proteins build on significant literature using classical methods to model protein sequence variation. Statistical models of protein multiple sequence alignments, equivalent to Potts models from statistical physics, have been particularly widely used [7, 41–44]. Pairwise interaction terms from these models accurately predict crystal structure contacts, and these models are also useful for sequence generation and protein fitness prediction. Anishchenko *et al*. [12] previously used these statistical models to analyze how effects such as oligomeric contacts, structural differences in homologs, and alternative conformations impact protein evolution using statistical models, finding relatively small contributions from alternate conformations. Our analysis using the more powerful MSA Transformer model and molecular dynamics simulation data finds a significantly larger role of protein dynamics in residue-residue coevolution. These results demonstrate the utility of unsupervised deep learning models of protein sequence variation for scientific discovery, adding to work using such models for static structure prediction, sequence generation, and protein fitness prediction [8, 23, 45].

## Acknowledgements

This research is supported in part by an NIH grant R35-GM134922 (to YSS). We thank Naomi Latorraca for technical and general advice for processing and interpreting molecular dynamics simulation data. Antoine Koehl acknowledges the Miller Institute for Basic Research in Science for funding support.

## Supplementary Material

### A Supplementary Figures and Tables

**Fig. S1.**
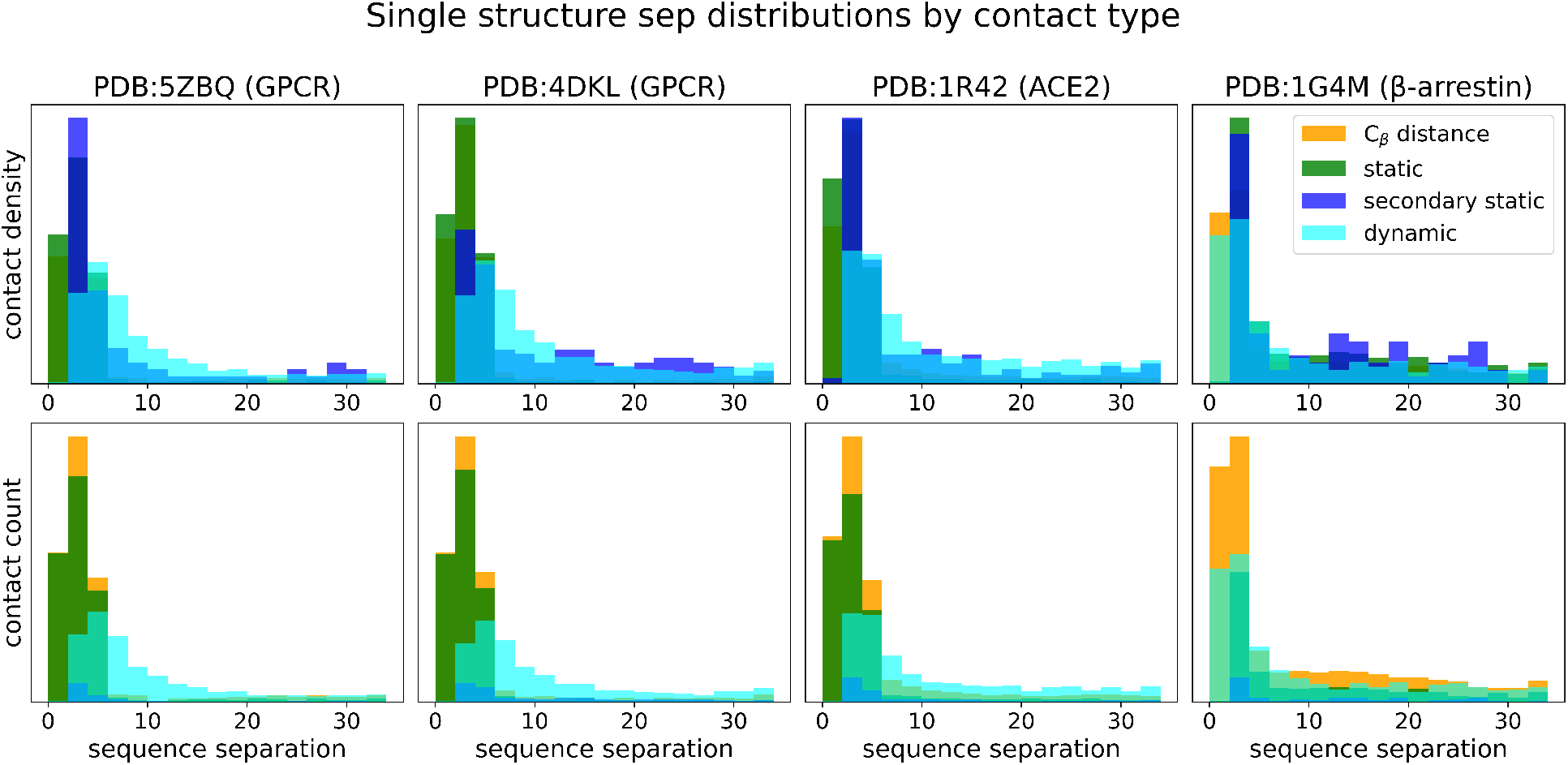
Residue-residue sequence separation distribution for unique contacts by contact type.

**Fig. S2.**
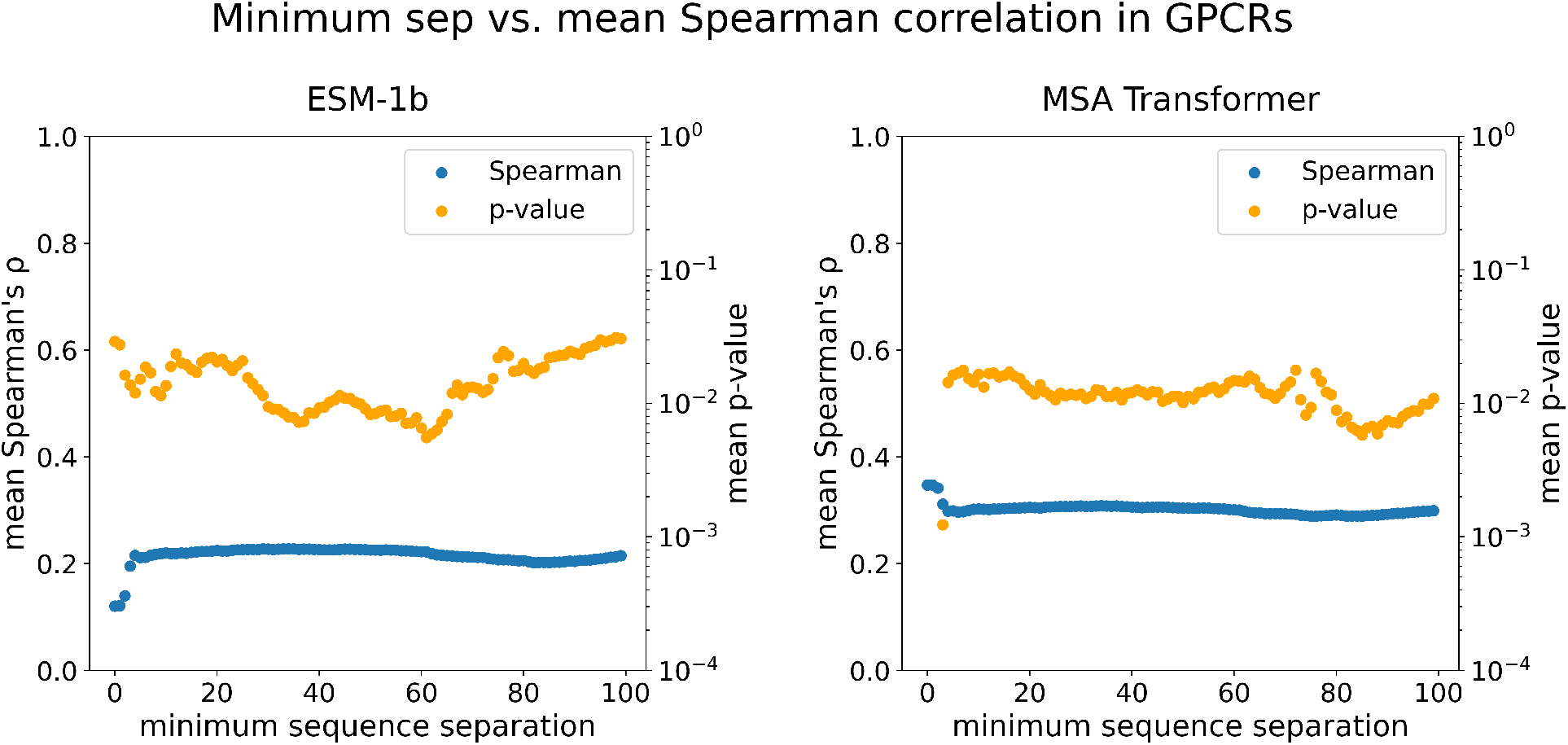
Mean Spearman’s rank correlation over all GPCRs by residue-residue sequence separation.

**Table S1.**
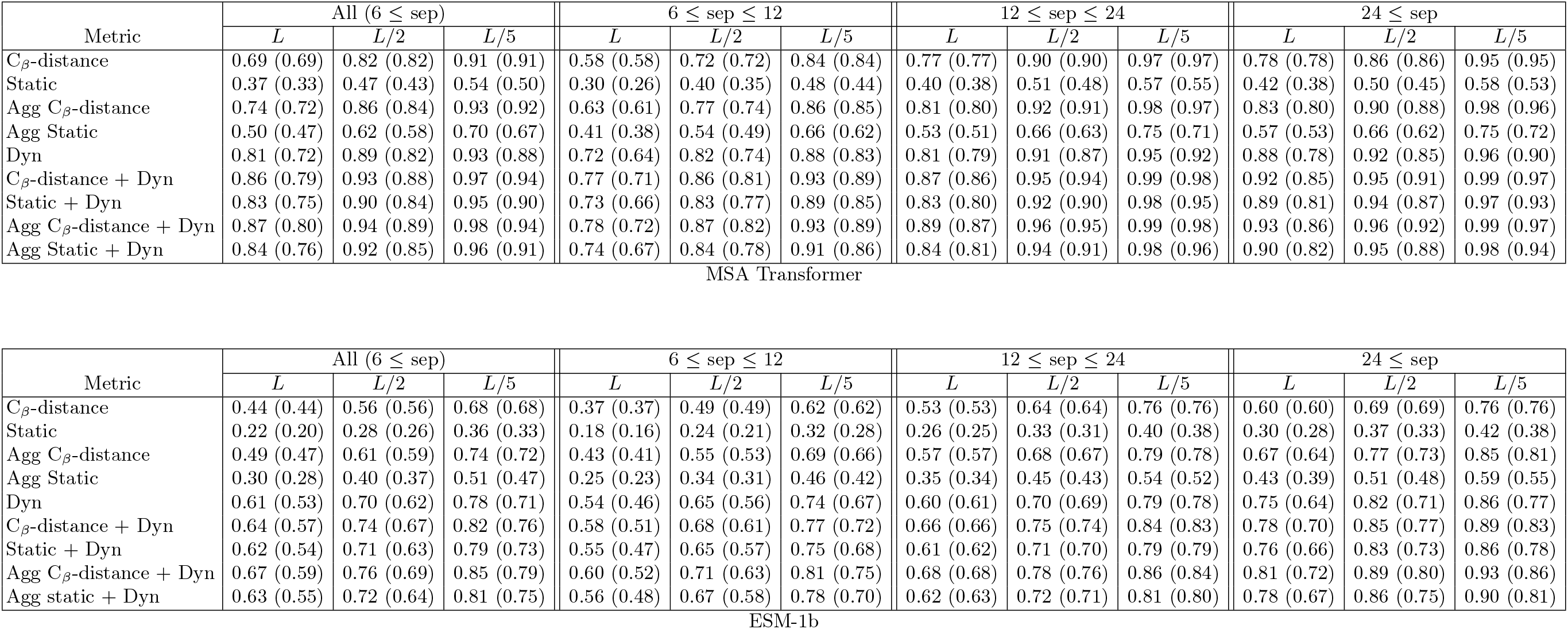
Full precision statistics for ESM-1b and MSA Transformer.

## References

1. D Altschuh, A M Lesk, A C Bloomer, and A Klug. Correlation of co-ordinated amino acid substitutions with function in viruses related to tobacco mosaic virus. J. Mol. Biol., 193(4):693–707, February 1987.

2. D Altschuh, T Vernet, P Berti, D Moras, and K Nagai. Coordinated amino acid changes in homologous protein families. Protein Eng., 2(3):193–199, September 1988.

3. Christopher S Miller and David Eisenberg. Using inferred residue contacts to distinguish between correct and incorrect protein models. Bioinformatics, 24(14):1575–1582, May 2008.

4. John Jumper, Richard Evans, Alexander Pritzel, Tim Green, Michael Figurnov, Olaf Ronneberger, Kathryn Tunyasuvunakool, Russ Bates, Augustin Žídek, Anna Potapenko, Alex Bridgland, Clemens Meyer, Simon A A Kohl, Andrew J Ballard, Andrew Cowie, Bernardino Romera-Paredes, Stanislav Nikolov, Rishub Jain, Jonas Adler, Trevor Back, Stig Petersen, David Reiman, Ellen Clancy, Michal Zielinski, Martin Steinegger, Michalina Pacholska, Tamas Berghammer, Sebastian Bodenstein, David Silver, Oriol Vinyals, Andrew W Senior, Koray Kavukcuoglu, Pushmeet Kohli, and Demis Hassabis. Highly accurate protein structure prediction with AlphaFold. Nature, 596(7873):583–589, July 2021.

5. Minkyung Baek, Frank DiMaio, Ivan Anishchenko, Justas Dauparas, Sergey Ovchinnikov, Gyu Rie Lee, Jue Wang, Qian Cong, Lisa N Kinch, R Dustin Schaeffer, et al. Accurate prediction of protein structures and interactions using a three-track neural network. Science, 373(6557):871–876, 2021.

6. Zeming Lin, Halil Akin, Roshan Rao, Brian Hie, Zhongkai Zhu, Wenting Lu, Allan dos Santos Costa, Maryam Fazel-Zarandi, Tom Sercu, Sal Candido, et al. Language models of protein sequences at the scale of evolution enable accurate structure prediction. bioRxiv, 2022.

7. Debora S Marks, Lucy J Colwell, Robert Sheridan, Thomas A Hopf, Andrea Pagnani, Riccardo Zecchina, and Chris Sander. Protein 3D structure computed from evolutionary sequence variation. PLoS One, 6(12):e28766, December 2011.

8. Roshan Rao, Joshua Meier, Tom Sercu, Sergey Ovchinnikov, and Alexander Rives. Transformer protein language models are unsupervised structure learners. In Proceedings of International Conference on Learning Representations, 2021.

9. Kathryn Tunyasuvunakool, Jonas Adler, Zachary Wu, Tim Green, Michal Zielinski, Augustin Žídek, Alex Bridg-land, Andrew Cowie, Clemens Meyer, Agata Laydon, et al. Highly accurate protein structure prediction for the human proteome. Nature, 596(7873):590–596, 2021.

10. Diego Del Alamo, Davide Sala, Hassane S Mchaourab, and Jens Meiler. Sampling alternative conformational states of transporters and receptors with alphafold2. eLife, 11:e75751, 2022.

11. Scott A Hollingsworth and Ron O Dror. Molecular dynamics simulation for all. Neuron, 99(6):1129–1143, 2018.

12. Ivan Anishchenko, Sergey Ovchinnikov, Hetunandan Kamisetty, and David Baker. Origins of coevolution between residues distant in protein 3D structures. Proceedings of the National Academy of Sciences, 114(34):9122–9127, 2017.

13. Sarel J Fleishman and Amnon Horovitz. Extending the new generation of structure predictors to account for dynamics and allostery. J. Mol. Biol., 433(20):167007, October 2021.

14. Roshan M Rao, Jason Liu, Robert Verkuil, Joshua Meier, John Canny, Pieter Abbeel, Tom Sercu, and Alexander Rives. MSA transformer. In Marina Meila and Tong Zhang, editors, Proceedings of the 38th International Conference on Machine Learning, volume 139 of Proceedings of Machine Learning Research, pages 8844–8856. PMLR, 2021.

15. Bulat Faezov and Roland L Dunbrack, Jr. PDBrenum: A webserver and program providing protein data bank files renumbered according to their UniProt sequences. PLoS One, 16(7):e0253411, July 2021.

16. Helen M Berman, John Westbrook, Zukang Feng, Gary Gilliland, T N Bhat, Helge Weissig, Ilya N Shindyalov, and Philip E Bourne. The protein data bank. Nucleic Acids Res., 28(1):235–242, January 2000.

17. UniProt Consortium. UniProt: the universal protein knowledgebase in 2021. Nucleic Acids Res., 49(D1):D480–D489, January 2021.

18. Ismael Rodríguez-Espigares, Mariona Torrens-Fontanals, Johanna K S Tiemann, David Aranda-García, Juan Manuel Ramírez-Anguita, Tomasz Maciej Stepniewski, Nathalie Worp, Alejandro Varela-Rial, Adrián Morales-Pastor, Brian Medel-Lacruz, Gáspár Pándy-Szekeres, Eduardo Mayol, Toni Giorgino, Jens Carlsson, Xavier Deupi, Slawomir Filipek, Marta Filizola, José Carlos Gómez-Tamayo, Angel Gonzalez, Hugo Gutiérrez-de Terán, Mireia Jiménez-Rosés, Willem Jespers, Jon Kapla, George Khelashvili, Peter Kolb, Dorota Latek, Maria Marti-Solano, Pierre Matricon, Minos-Timotheos Matsoukas, Przemyslaw Miszta, Mireia Olivella, Laura Perez-Benito, Davide Provasi, Santiago Ríos, Iván R Torrecillas, Jessica Sallander, Agnieszka Sztyler, Silvana Vasile, Harel Weinstein, Ulrich Zachariae, Peter W Hildebrand, Gianni De Fabritiis, Ferran Sanz, David E Gloriam, Arnau Cordomi, Ramon Guixà-González, and Jana Selent. GPCRmd uncovers the dynamics of the 3D-GPCRome. Nat. Methods, 17(8):777–787, August 2020.

19. Naomi R Latorraca, Matthieu Masureel, Scott A Hollingsworth, Franziska M Heydenreich, Carl-Mikael Suomivuori, Connor Brinton, Raphael J L Townshend, Michel Bouvier, Brian K Kobilka, and Ron O Dror. How GPCR phosphorylation patterns orchestrate Arrestin-Mediated signaling. Cell, 183(7):1813–1825.e18, December 2020.

20. D.E. Shaw Research. Molecular Dynamics Simulations Related to SARS-CoV-2. D.E. Shaw Research Technical Data https://www.deshawresearch.com/downloads/download_trajectory_sarscov2.cgi, 2020.

21. Fonseca, Rasumus et al. GetContacts. https://getcontacts.github.io/, 2022.

22. Jack Holland, Qinxin Pan, and Gevorg Grigoryan. Contact prediction is hardest for the most informative contacts, but improves with the incorporation of contact potentials. PLoS One, 13(6):e0199585, June 2018.

23. Alexander Rives, Joshua Meier, Tom Sercu, Siddharth Goyal, Zeming Lin, Jason Liu, Demi Guo, Myle Ott, C Lawrence Zitnick, Jerry Ma, and Rob Fergus. Biological structure and function emerge from scaling unsupervised learning to 250 million protein sequences. Proceedings of the National Academy of Sciences, 118(15):e2016239118, 2021.

24. Michael Remmert, Andreas Biegert, Andreas Hauser, and Johannes Söding. HHblits: lightning-fast iterative protein sequence searching by HMM-HMM alignment. Nature Methods, 9(2):173–175, December 2011.

25. Ashish Vaswani, Noam Shazeer, Niki Parmar, Jakob Uszkoreit, Llion Jones, Aidan N Gomez, L Ukasz Kaiser, and Illia Polosukhin. Attention is all you need. In I Guyon, U Von Luxburg, S Bengio, H Wallach, R Fergus, S Vishwanathan, and R Garnett, editors, Advances in Neural Information Processing Systems, volume 30. Curran Associates, Inc., 2017.

26. Jesse Vig, Ali Madani, Lav R Varshney, Caiming Xiong, Richard Socher, and Nazneen Fatema Rajani. BERTology meets biology: Interpreting attention in protein language models. bioRxiv preprint: 2020.06.26.174417, June 2020.

27. William R Taylor and Michael I Sadowski. Structural constraints on the covariance matrix derived from multiple aligned protein sequences. PLoS One, 6(12):e28265, December 2011.

28. Hetunandan Kamisetty, Sergey Ovchinnikov, and David Baker. Assessing the utility of coevolution-based residue–residue contact predictions in a sequence- and structure-rich era. Proceedings of the National Academy of Sciences, 110(39):15674–15679, 2013.

29. Iakes Ezkurdia, Osvaldo Graña, José M G Izarzugaza, and Michael L Tress. Assessment of domain boundary predictions and the prediction of intramolecular contacts in CASP8. Proteins, 77 Suppl 9:196–209, 2009.

30. Thóra K Bjarnadóttir, David E Gloriam, Sofia H Hellstrand, Helena Kristiansson, Robert Fredriksson, and Helgi B Schiöth. Comprehensive repertoire and phylogenetic analysis of the G protein-coupled receptors in human and mouse. Genomics, 88(3):263–273, September 2006.

31. Daniel Hilger, Matthieu Masureel, and Brian K Kobilka. Structure and dynamics of GPCR signaling complexes. Nat. Struct. Mol. Biol., 25(1):4–12, January 2018.

32. William I Weis and Brian K Kobilka. The molecular basis of G Protein-Coupled receptor activation. Annu. Rev. Biochem., 87:897–919, June 2018.

33. Søren G F Rasmussen, Hee-Jung Choi, Juan Jose Fung, Els Pardon, Paola Casarosa, Pil Seok Chae, Brian T Devree, Daniel M Rosenbaum, Foon Sun Thian, Tong Sun Kobilka, Andreas Schnapp, Ingo Konetzki, Roger K Sunahara, Samuel H Gellman, Alexander Pautsch, Jan Steyaert, William I Weis, and Brian K Kobilka. Structure of a nanobody-stabilized active state of the *β*(2) adrenoceptor. Nature, 469(7329):175–180, January 2011.

34. Daniel M Rosenbaum, Cheng Zhang, Joseph A Lyons, Ralph Holl, David Aragao, Daniel H Arlow, Søren G F Rasmussen, Hee-Jung Choi, Brian T Devree, Roger K Sunahara, Pil Seok Chae, Samuel H Gellman, Ron O Dror, David E Shaw, William I Weis, Martin Caffrey, Peter Gmeiner, and Brian K Kobilka. Structure and function of an irreversible agonist-*β*(2) adrenoceptor complex. Nature, 469(7329):236–240, January 2011.

35. Aashish Manglik, Tae Hun Kim, Matthieu Masureel, Christian Altenbach, Zhongyu Yang, Daniel Hilger, Michael T Lerch, Tong Sun Kobilka, Foon Sun Thian, Wayne L Hubbell, R Scott Prosser, and Brian K Kobilka. Structural insights into the dynamic process of *β*2-Adrenergic receptor signaling. Cell, 161(5):1101–1111, May 2015.

36. Fang Li, Wenhui Li, Michael Farzan, and Stephen C Harrison. Structure of SARS coronavirus spike receptor-binding domain complexed with receptor. Science, 309(5742):1864–1868, September 2005.

37. Paul Towler, Bart Staker, Sridhar G Prasad, Saurabh Menon, Jin Tang, Thomas Parsons, Dominic Ryan, Martin Fisher, David Williams, Natalie A Dales, Michael A Patane, and Michael W Pantoliano. ACE2 x-ray structures reveal a large hinge-bending motion important for inhibitor binding and catalysis. J. Biol. Chem., 279(17):17996–18007, April 2004.

38. Ellen D Zhong, Tristan Bepler, Bonnie Berger, and Joseph H Davis. CryoDRGN: reconstruction of heterogeneous cryo-EM structures using neural networks. Nat. Methods, 18(2):176–185, February 2021.

39. Ali Punjani and David Fleet. 3D flexible refinement: Structure and motion of flexible proteins from cryo-em. Microscopy and Microanalysis, 28(S1):1218–1218, 2022.

40. Dan Rosenbaum, Marta Garnelo, Michal Zielinski, Charlie Beattie, Ellen Clancy, Andrea Huber, Pushmeet Kohli, Andrew W Senior, John Jumper, Carl Doersch, S M Ali Eslami, Olaf Ronneberger, and Jonas Adler. Inferring a continuous distribution of atom coordinates from Cryo-EM images using VAEs. arXiv preprint arXiv:2106.14108, June 2021.

41. Magnus Ekeberg, Cecilia Lövkvist, Yueheng Lan, Martin Weigt, and Erik Aurell. Improved contact prediction in proteins: Using pseudolikelihoods to infer potts models. Phys. Rev. E, 87(1):012707, January 2013.

42. Martin Weigt, Robert A White, Hendrik Szurmant, James A Hoch, and Terence Hwa. Identification of direct residue contacts in protein–protein interaction by message passing. Proceedings of the National Academy of Sciences, 106(1):67–72, 2009.

43. R B Potts. Some generalized order-disorder transformations. Math. Proc. Cambridge Philos. Soc., 48(1):106–109, January 1952.

44. Faruck Morcos, Andrea Pagnani, Bryan Lunt, Arianna Bertolino, Debora S Marks, Chris Sander, Riccardo Zecchina, José N Onuchic, Terence Hwa, and Martin Weigt. Direct-coupling analysis of residue coevolution captures native contacts across many protein families. Proc. Natl. Acad. Sci. U. S. A., 108(49):E1293–301, December 2011.

45. Adam J Riesselman, John B Ingraham, and Debora S Marks. Deep generative models of genetic variation capture the effects of mutations. Nature Methods, 15(10):816–822, 2018.

